## Supplementary Information for "Discovery of a non-canonical prototype long-chain monoacylglycerol lipase through a structure-based endogenous reaction intermediate complex"

### Supplementary Figures

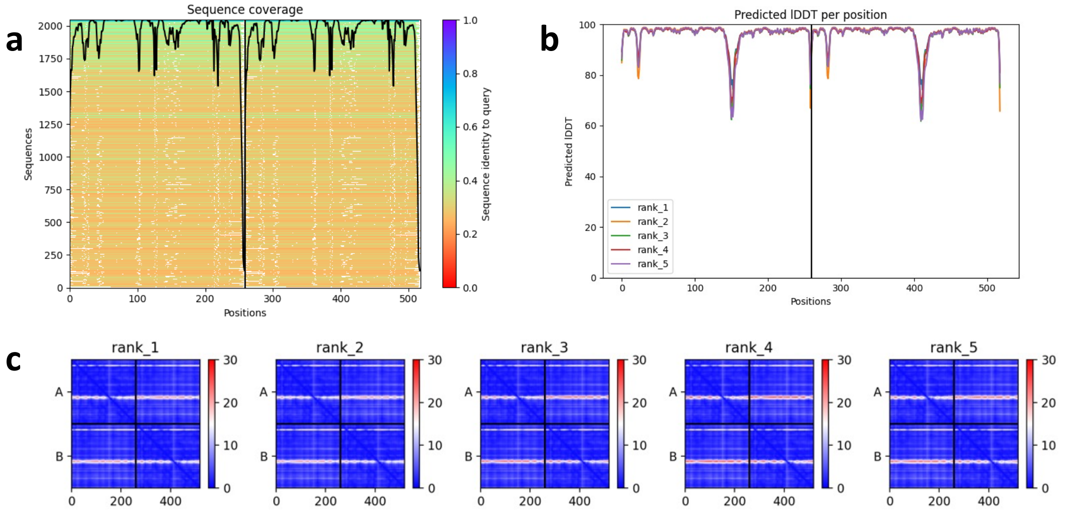

**Supplementary Figure 1. Alpha Fold 2 predicted structures of dimeric *Tth*** MAG lipase The overall root mean square deviations of the five top predicted structures with the experimental *Tth* MAG lipase structures are in the range of 0.4-0.5 Å. The majority of the *Tth* dimer interface specific interactions **(Fig. 2c)** are not recovered in the predicted structures. In addition, any information on protein-ligand interactions (**Fig. 3**) is missing in the predicted models. The vertical and horizontal lines shown in the plots indicate the border between the first and second sequence chains (A, B) of the predicted models. **a**, Sequence coverage of the predicted models; **b,** Local Distance Difference Test, indicating a major peak within the sequence segment of the *Tth* MAG lipase lid domain; **c,** Distance matrices of the top five ranking models, indicating the highest level of uncertainty at the sequence segment of the *Tth* MAG lipase lid domain.

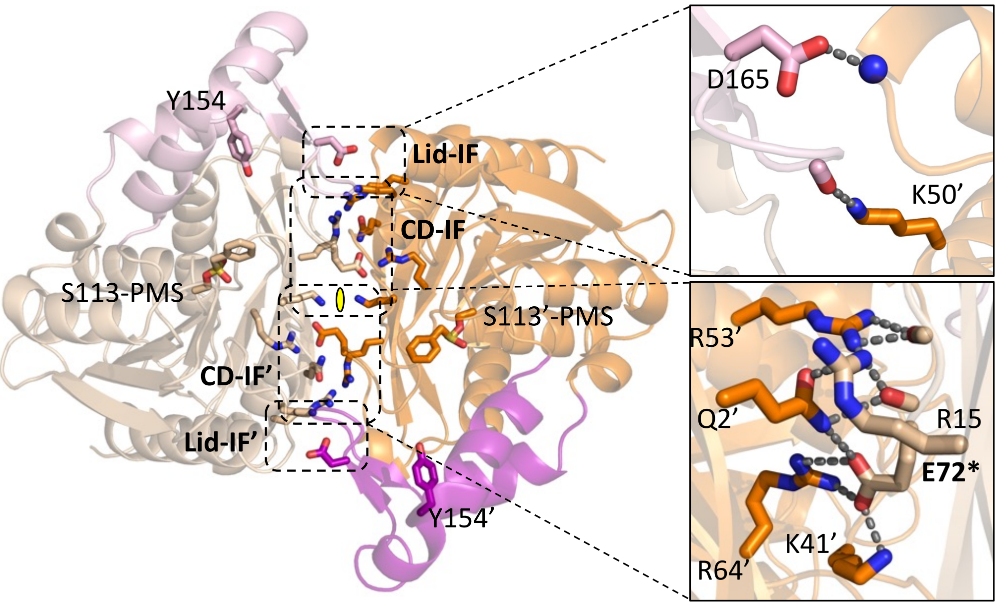

**Supplementary Figure 2. Two-fold symmetric dimeric interface of *Tth* MAG lipase. T**wo-fold symmetry is indicated by a yellow symbol. The majority of specific interactions originate from the two-fold repeated central CD interface labeled CD-IF and CD-IF', and a smaller number of interactions originate from the two-fold repeated distal lid domain interface labeled Lid-IF and Lid-IF'. Zoom-in of the interfaces are shown on the right indicating specific hydrogen bonds (cf. panel D, Supplementary Figure S2). As the PMSF-inhibited *Tth* MAG lipase structure is shown, Ser113-PMS is labeled in both protomers, to approximately locate the two active sites. The position of Tyr154, which was used for mutational analysis, is also labeled. Each *Tth* MAG lipase molecule comprises a two-domain structure. The larger CD is assembled by two sequence segments (residues 1-140 and 184-259), which form a well-established α/β-hydrolase fold comprising a central eight-stranded superhelical twisted β-sheet and six α-helices positioned on both sides of the sheet. In the PMSF-inhibited *Tth* MAG lipase structure, a covalent PMS-adduct with the catalytic Ser113 is formed, thus blocking the active site from further enzymatic reaction ligands (**Fig. 3B**). In agreement with other lipase structures, *Tth* MAG lipase Ser113 is located at the nucleophilic elbow directly after strand β5 to act as a catalytic residue adopting a strained ε-conformation **(Fig. 3a-c, Supplementary Fig. S3**). This position is part of the catalytic triad, which also includes Asp203 and His233 **(Fig. 3a-b, Supplementary Fig. 2).** In both structures an active oxyanion hole is formed by Phe37 and Met114 by orienting their main chain amido groups towards the MAG lipase active site **(Fig. 3b).**

**a**

**b**

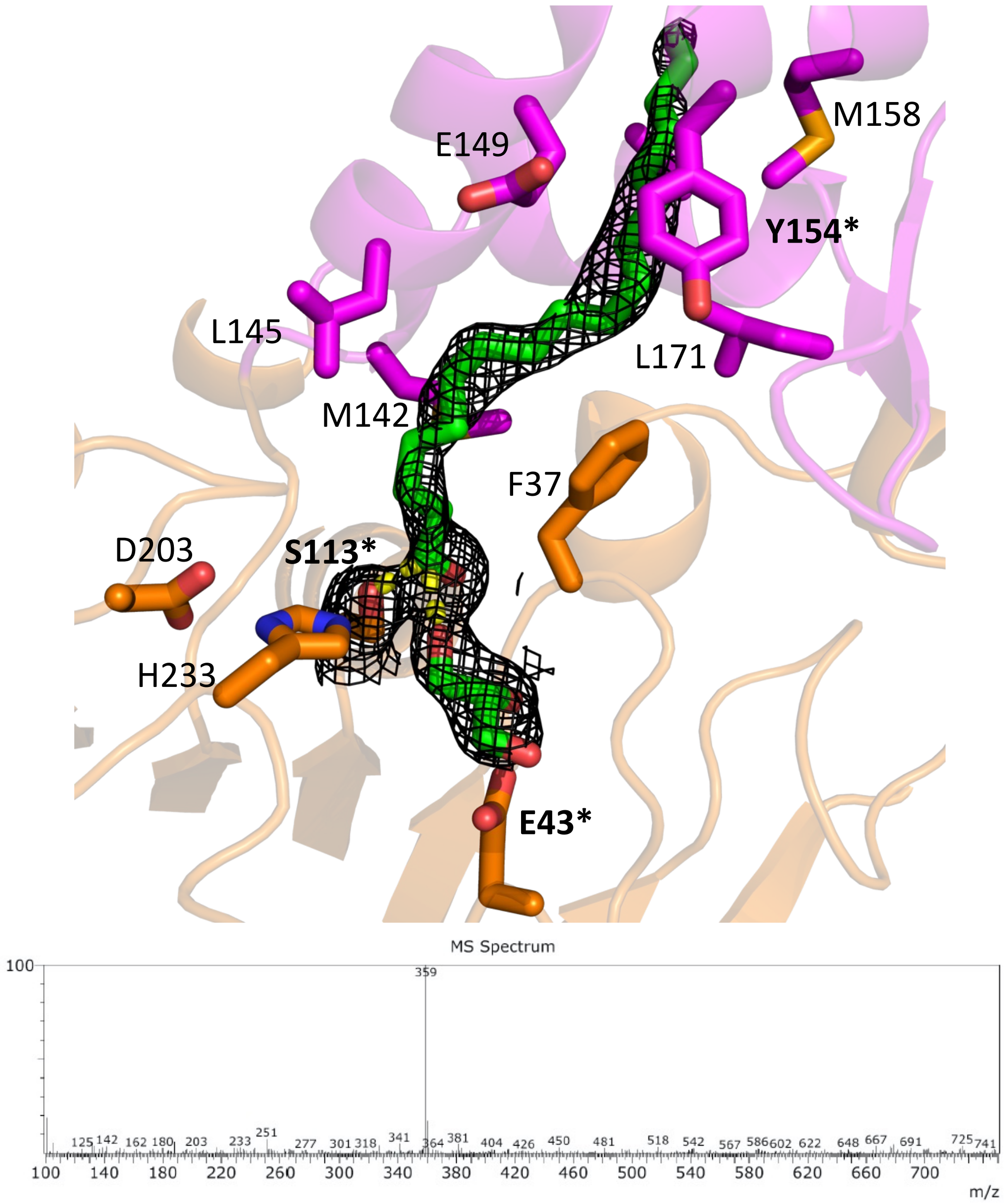

**Supplementary Figure 3. a**, evidence for the presence of endogenous C18 MAG intermediate in the *Tth* MAG lipase structure without PMSF. *2Fo-Fc* omit electron density map at 1 σ contour level. *Tth* MAG lipase residues that are involved in binding to C18 MAG intermediate are shown and labeled (*cf*. **Fig. 3a**). **b,** reverse phase chromatography analysis of bound ligands in *Tth* MAG lipase crystals in the absence of PMSF**.** Source data are provided within **Supplementary Data 2.**

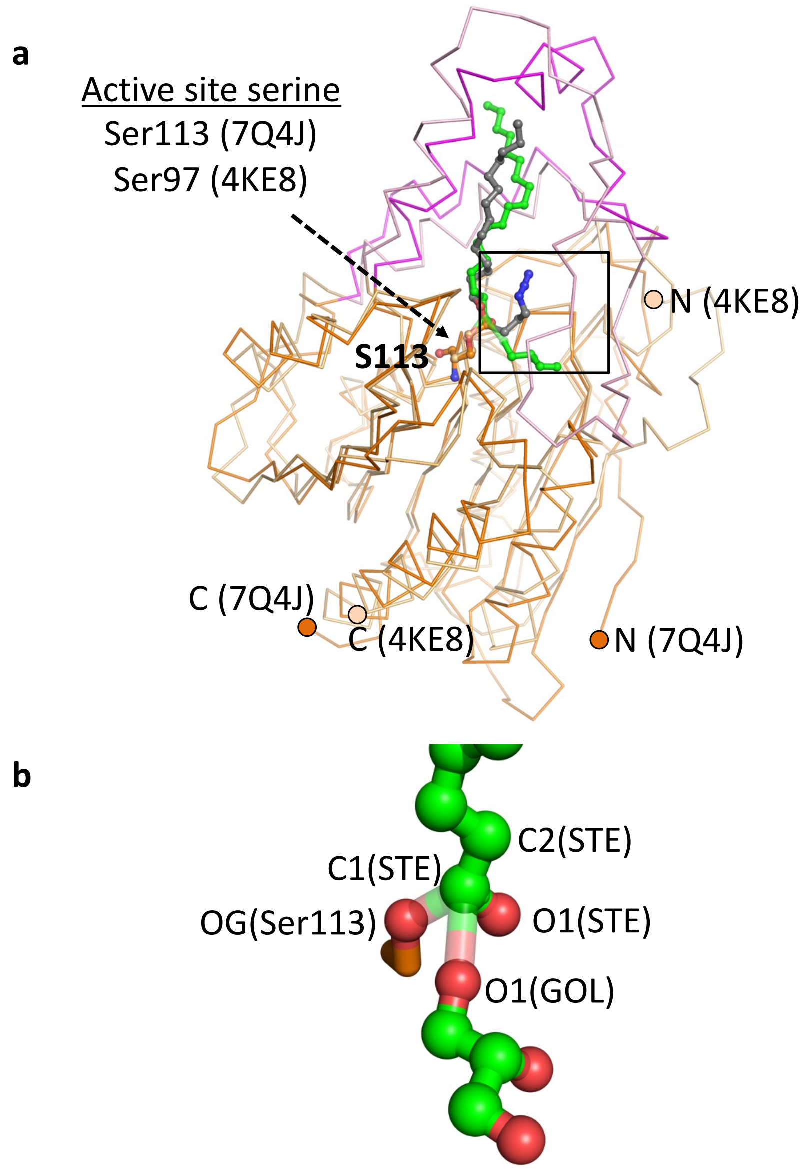

**Supplementary Figure 4. Geometry of MAG intermediate in *Tth* MAG lipase. a**, superposition of *Tth* MAG lipase in complex with C18 MAG intermediate and *Bacillus sp.* MAG lipase in complex with C14 MAG analogue (PDB entry: 4KE8). Colors for the *Tth* MAG lipase complex are as in **Fig. 1** and **2.** Colors for *Bacillus sp.* MAG lipase in complex with C14 MAG analogue: protein, light orange; MAG analogue, grey. Oxygen and nitrogen atoms are in red and blue, respectively. **b**, zoom into *Tth* MAG lipase MAG tetrahedral intermediate. Transient bonds are in pale colors. Atoms are specified as in **Supplementary Table 11.**

**a**

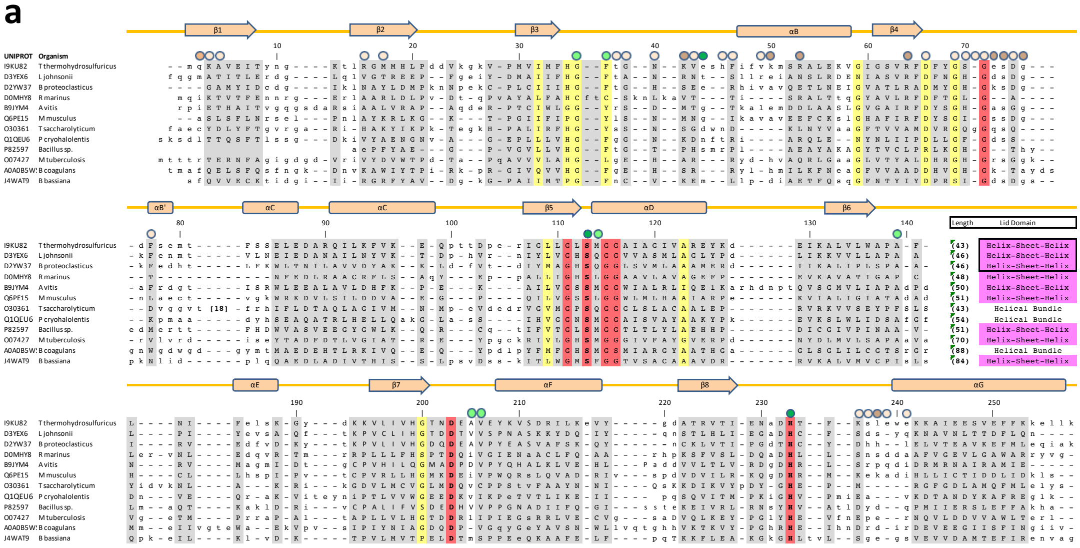

**b**

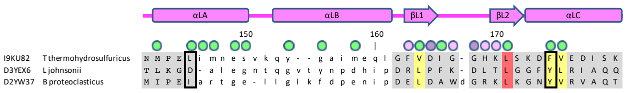

**Supplementary Figure 5. a**, alignment of 12 CD sequences, for which an overall structure-based alignment was possible. UNIPROT codes and organisms of aligned sequences are indicated to the left. Structurally aligned residues are shown in capital letters and respective aligned positions are shaded in grey; other residues are in small letters. As a structure-based alignment of the lid domain segment, inserted into the N-terminal and C-terminal part of each CD sequence, was not possible due to high level of sequence and structural diversity, instead the length and type of lid domains of all chosen esterase/lipases are indicated. HBH-type lid domains with structural relations to the *Tth* MAG lipase lid domain are colored in magenta. Sequence numbers indicated in 10-residues intervals on top of the alignment refer to the *Tth* MAG lipase sequence. Helices and β-strands, indicated by cylinders and arrows, respectively, are taken from the *Tth* MAG lipase coordinates. The coloring codes of these secondary structural elements are as in **Fig. 2**. Long insertions of single sequences are shown by numbers, indicating the length of the inserted sequence. Conserved residue positions are highlighted in red (invariant) or yellow (highly conserved). Residues involved in ligand interactions (C18 MAG) in the *Tth* MAG lipase structure without PMSF are indicated by green circles on top of the alignment: hydrogen bond interactions, dark green; other interactions, light green. Residues involved in *Tth* MAG lipase dimer interactions are indicated by orange circles: hydrogen bond interactions, dark orange; other interactions, light orange. **b,** alignment of the HBH lid domains of three two sequences closely related to *Tth* MAG lipase, for which a structure-based alignment was possible. The color codes are as for panel a, except residues involved into lid domain dimer interface interactions are highlighted with magenta circles according to the color codes from **Fig. 2** on top of the alignment. Residues with changed properties for active site ligand binding are boxed.

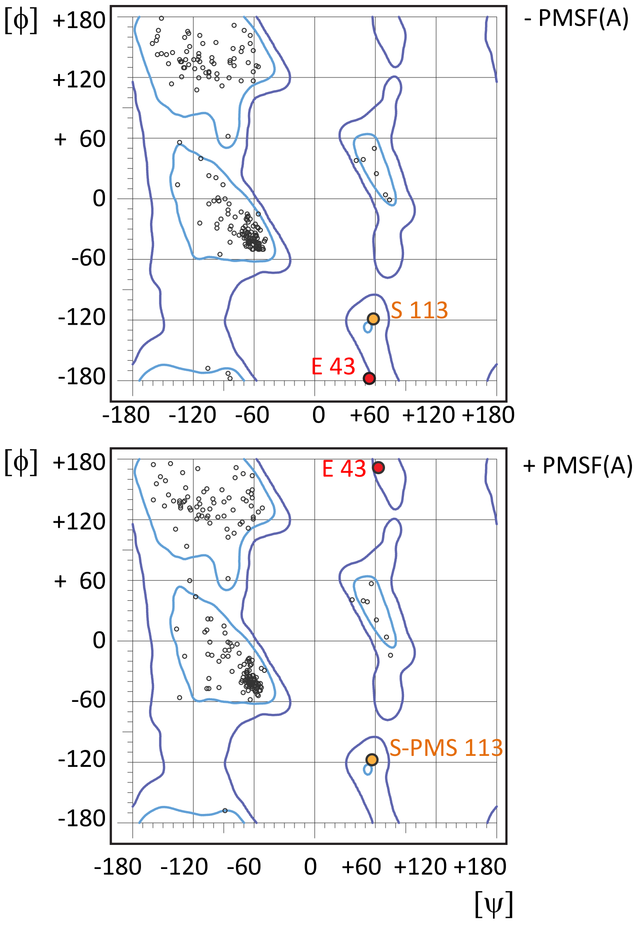

**Supplementary Figure 6.** Ramachandran plot of *Tth* MAG lipase structures, in the absence and presence of PMSF. In both structures, Glu43 and Ser113 have unusual dihedral angles, due to functional reasons (for details, see text).

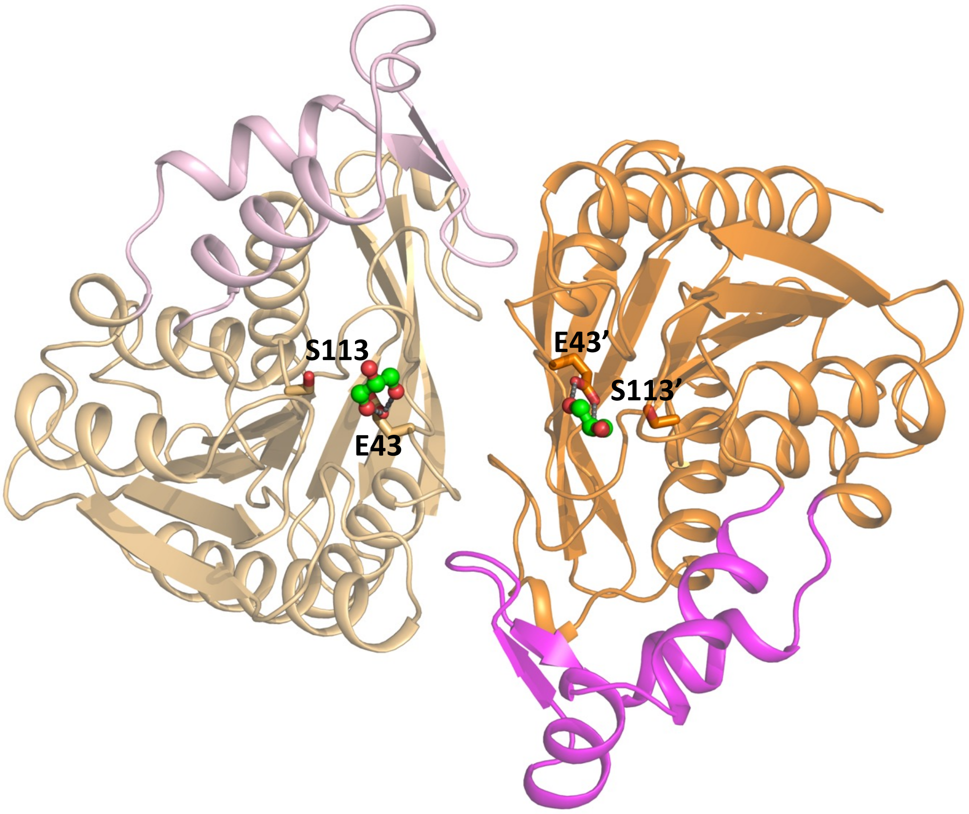

**Supplementary Figure 7.** Structure of native, non-methylated *Tth* MAG lipase (PDB code: 8B9S), demonstrating the presence of glycerol in the *Tth* MAG lipase active site. Color codes are as in **Fig. 1** and **2.** In contrast to the structure of methylated *Tth* MAG lipase (PDB entry: 7Q4J) there is no evidence for an acylglyeride intermediate.

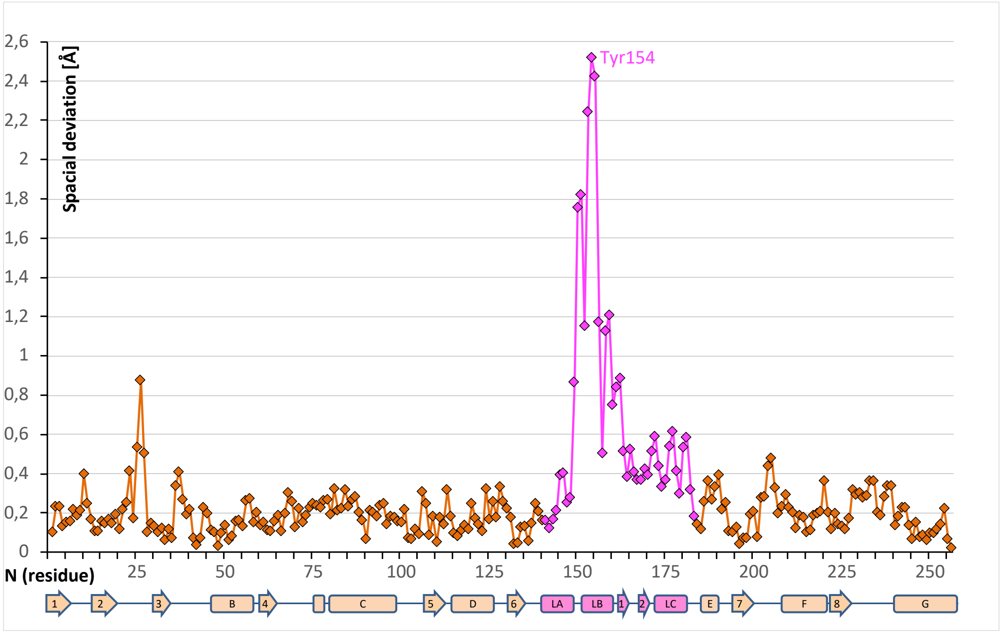

**Supplementary Figure 8. Dynamic substrate tunnel opening in the *Tth* MAG lipase lid domain** (*cf*. **Fig. 3D-E**). Spatial differences of the Cα positions of the *Tth* MAG lipase structures, with and without PMSF. Color codes are as in **Fig. 2.** The approximate positions of secondary structural elements are indicated **(Fig. 2a, Supplementary Fig. 2)**. The comparison indicates peak movements of lid domain residues next to the tunnel opener Tyr154.

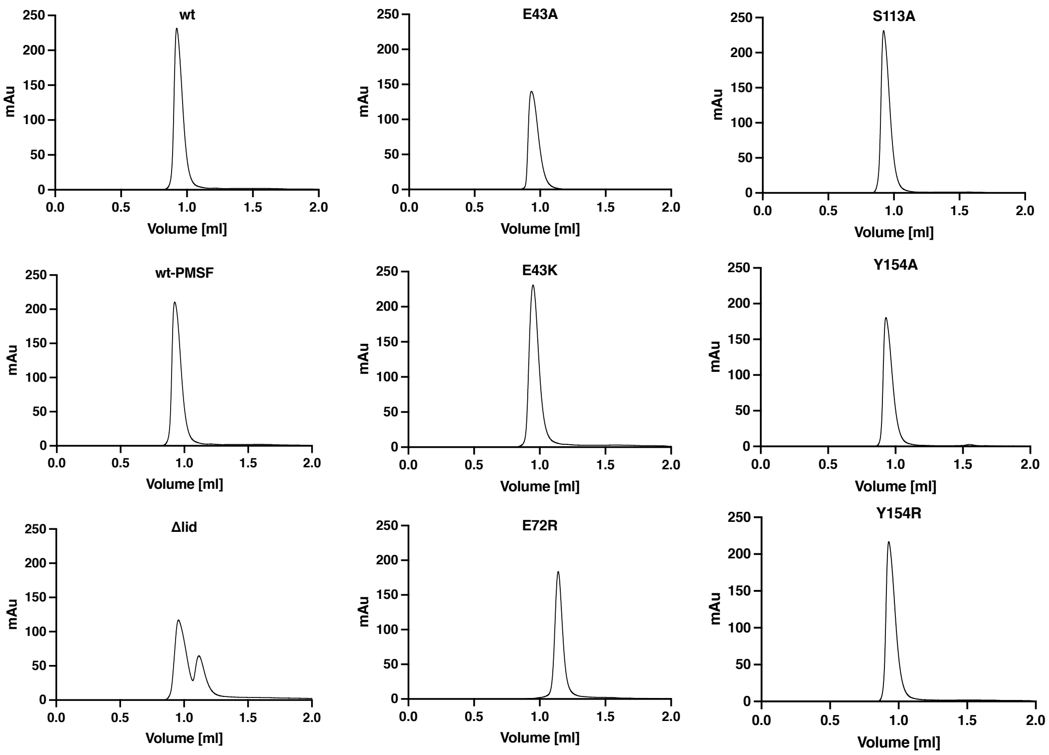

**Supplementary Figure 9. Analytic SEC profiles of all *Tth* MAG lipase variants investigated.** The respective elution profiles – except the ΔLid and E72R mutants – were interpreted as showing a dimeric state of the enzyme. Detection has been in milli Absorbance Units [mAU]. Source data are provided as a Source Data file.

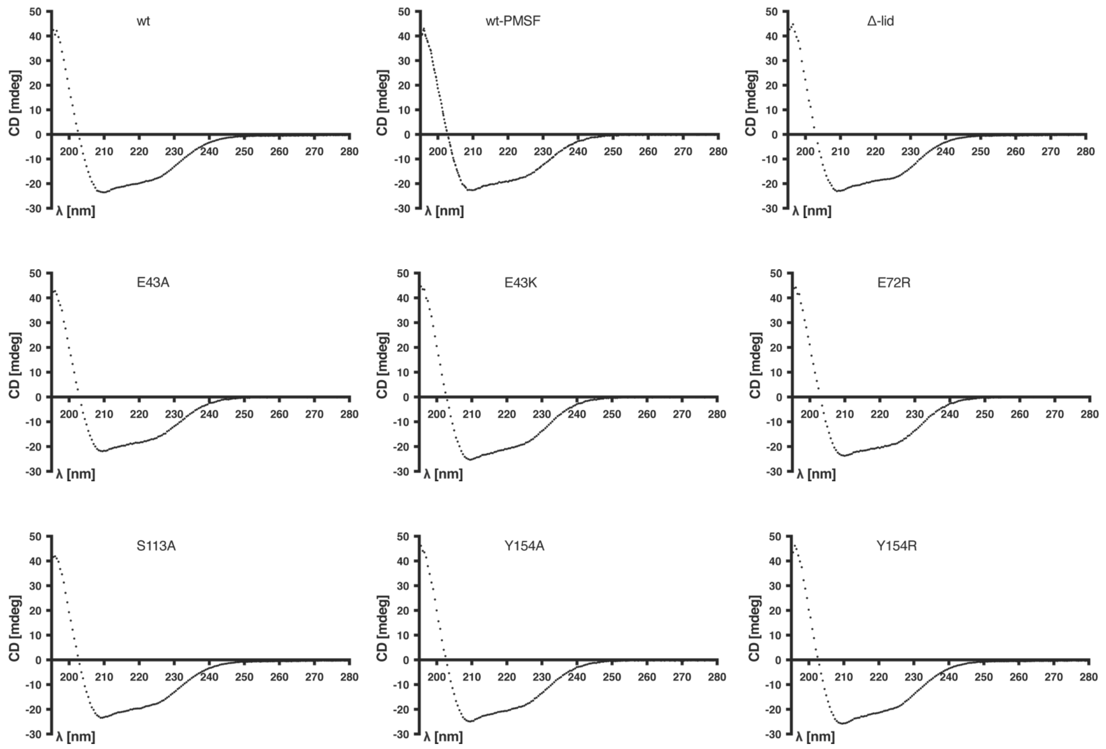

**Supplementary Figure 10.** CD spectra for all MAG lipase *Tth* variants investigated, showing no significant deviations from wt *Tth* MAG lipase. Samples were measured in triplicates. For quantification of secondary structure content, see **Supplementary Table 4.** Source data are provided as a Source Data file.

**a**

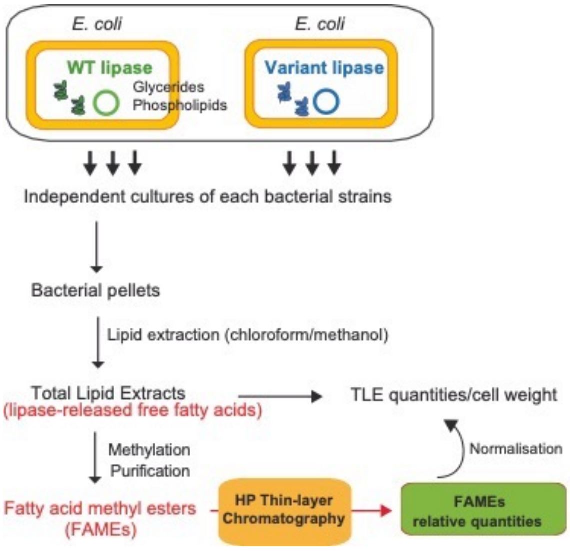

**b**

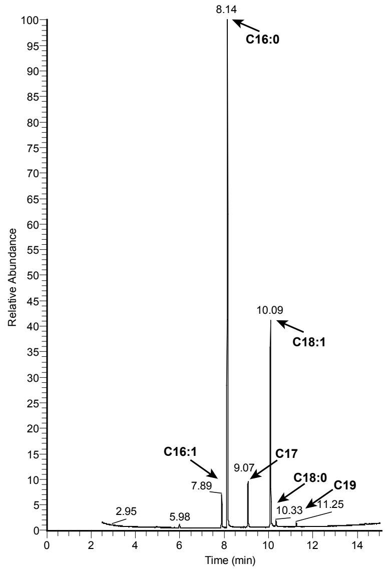

**Supplementary Figure 11. a,** Schematic workflow of sample preparation and lipid cell-based data analysis. *E. coli* cultures were transformed with *Tth* MAG lipase variants, induced with IPTG for 60 min. Bacterial pellets were used to prepare total lipid extracts (TLE) following chloroform/methanol extraction. Lipid extracts were methylated to yield Fatty Acid Methyl Esters (FAMEs), further purified though florisil columns and the resulting enriched FAMEs were analyzed by high-pressure thin-layer chromatography (HPTLC) and quantified. Plate visualization was realized using primuline dye and scanning. Fluorescence associated with FAME bands was quantified, and normalization to TLE and bacterial mass was performed to correct for variations across samples. **b,** representative gas chromatogram used for FAME analysis.

a

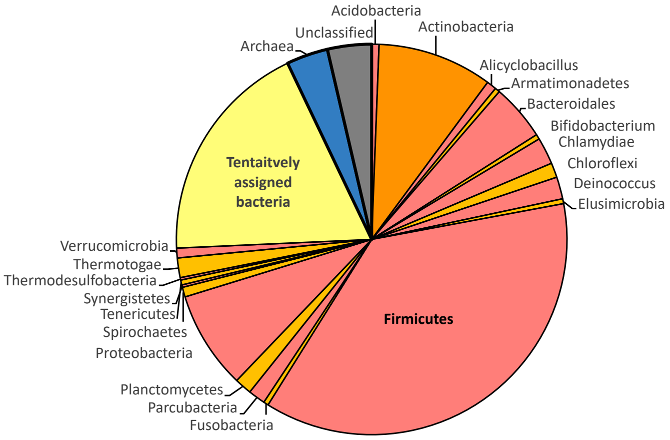

.

b

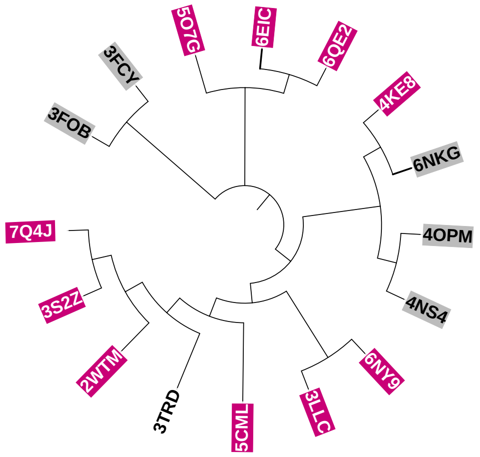

**Supplementary Figure 12. a,** taxonomic analysis of most closely related 496 sequence groups as defined in UniprotKB50, indicating that all related sequences (to the extent unambiguously assigned) are either of bacterial or archaeal origin. The statistics for different bacterial phyla are shown in alphabetic order and in alternating colors (red, orange). Statistics of unclassified or tentatively assigned bacterial sequences are shown in yellow, archaeal sequences in blue, and unclassified sequences are in grey. The complete list of sequence groups is in **Supplementary Data 1**. **b,** phylogenetic tree presentation of sequences with known 3D structures and distant relations to *Tth* MAG lipase, indicating the respective PDB ID codes (*cf***. Supplementary Table 8**). Sequences containing a HBH lid domain are highlighted in pink **(*cf*. Fig. 7),** other sequences with helical lid domains are in grey. The precise phylogenetic distances are not reliable because of only distant relations and hence have no meaning in this presentation. Conditions for target selection: a) presence of a complete α/β hydrolase catalytic triad (Ser113, Asp203, His233 in the *Tth* MAG lipase sequence) to discard any false-positive hits with unrelated function, b) coverage of both the N-terminal and C-terminal segments of the *Tth* MAG lipase domain with the lid domain inserted in the resulting pairwise sequence alignments.

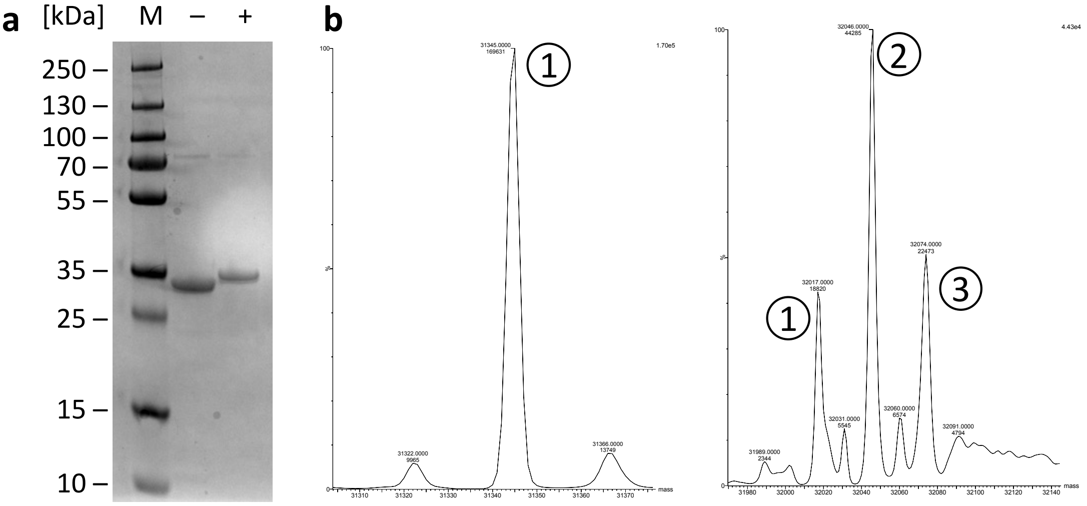

| Peak | Native *Tth* MGL [Da] | Methylated *Tth* MGL [Da] | Difference [Da] | # Methylation Sites |
|---|---|---|---|---|
| 1 | 31345 | 32017 | 672 | 24 |
| 2 | | 32046 | 701 | 25 |
| 3 | | 32074 | 729 | 26 |

**Supplementary Figure 13. Evidence for *Tth* MAG lipase methylation**. **a**, SDS-PAGE. M, marker; –, native *Tth* MAG lipase; +, methylated *Tth* MAG lipase. **b,** intact mass spectrometry: left, native *Tth* MAG lipase; right, methylated *Tth* MAG lipase. Measured molecular masses are listed below. The measured molecular mass of the native *Tth* MAG lipase includes the sequence of the N-terminal poly-histidine tag (MKHHHHHHSAGLEVLFQGP). Lysine di-methylation adds 28 Da per site, or when deprotonated 29 Da per site. The experiment was carried out once. Uncropped image of panel a) is provided in Source Data file. Source data from panel b) are provided within SourceDataPDF.zip file.

**a**

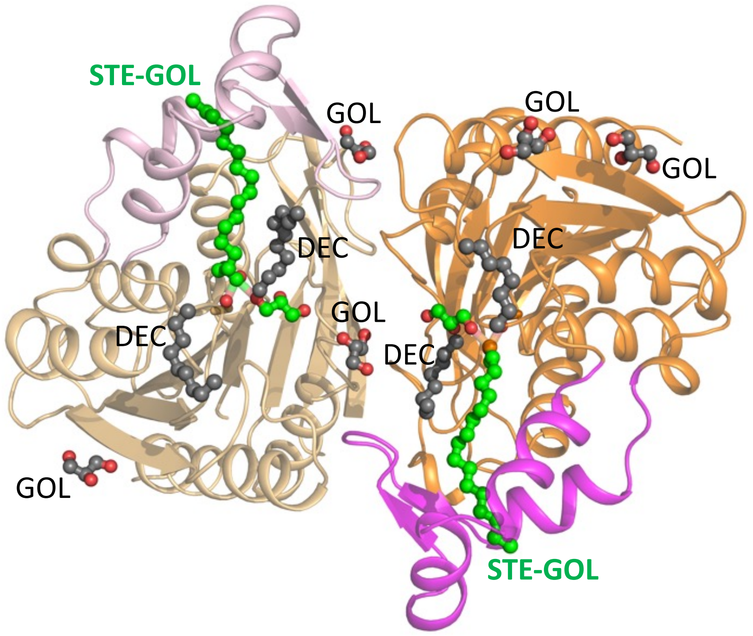

**b**

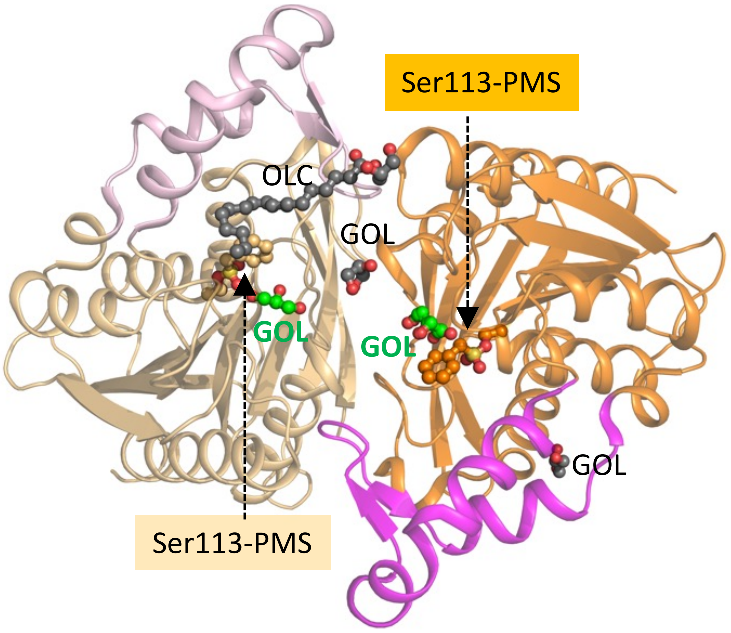

**Supplementary Figure 14.** Ligands found **a,** in the *Tth* MAG lipase Ser113-C18-MAG intermediate complex (PDB entry: 7Q4J) and **b,** in the *Tth* MAG lipase Ser113-PMS complex: STE-GOL, stearate-glycerol intermediate; DEC, alkane fragments tentatively assigned to contain 8,9 or 10 carbon atoms; OLC, 2,3-dihydroxypropyl-octadec-9-enoate; GOL, glycerol. Protein and validated ligand color codes are as in **Fig. 1** and **2**. Remaining, non-validated ligands are in grey and labeled.

### Supplementary Tables

**Supplementary Table 1: Thermostability of *Tth* variants**

| ***Tth* lipase variant** | **Tm (°C)** | **$\Delta T_m$(°C)** |
|---|---|---|
| wt | 77.2 ± 0.16 | |
| wt-PMS | 77.1 ± 0.04 | - 0.1 |
| ΔLid | 82.8 ± 0.01 | + 5.6 |
| E43A | 76.1 ± 0.02 | - 1.1 |
| E43K | 77.9 ± 0.21 | + 0.7 |
| E72R | 75.7 ± 0.05 | - 1.5 |
| S113A | 81.8 ± 0.07 | + 4.6 |
| Y154A | 78.3 ± 0.14 | + 1.1 |
| Y154R | 78.3 ± 0.10 | + 1.1 |

**Legend Supplementary Table 1:** We did not detect significant changes in thermostability (< 2 degrees changes), except for a variant without the lid domain (Δ140-183) that revealed an increased melting temperature of 83 °C, which is 6 °C higher than one found for the wt *Tth* MAG lipase **(Fig. 1b)**. This was within the melting temperature of the wt *Tth* MAG lipase **(Fig. 1b)**. This was within expectation as the lid domain is highly exposed and presents the most dynamic part of the overall *Tth* MAG lipase structure (**Fig. 1 & 2).** Interestingly, we have also observed a similar level of thermal stabilization in the *Tth* MAG lipase S113A active site mutant. The most plausible explanation is that a mutation of this active site residue could lead to a relieve of the high-energy ε-conformation of Ser113 found in both *wt* enzymes in the presence and absence of PMSF **(Supplementary Fig. 6)**. Data are presented as mean values ± SD, *n* = 3.

**Supplementary Table 2: Data collection, phasing and refinement statistics**

| | Native | Active Se-Met (methylated) | | PMS-inhibited (methylated) derivative |
|---|---|---|---|---|
| **Data collection** | | | | |
| Space group | $P3_2$ | $P4_32_12$ | | $P4_322$ |
| Cell dimensions | | | | |
| *a*, *b*, *c* (Å) | 125.2, 125.2, 101.4 | 137.5, 137.5, 67.8 | | 85.8, 85.8, 157.5 |
| $\alpha, \beta, \gamma$ (°) | 90, 90, 120 | 90, 90, 90 | | 90, 90, 90 |
| | | *Peak* | *Inflection* | |
| Wavelength | 0.9763 | 0.97873 | 0.97903 | 0.95372 |
| Resolution (Å) | 20.0 -2.42 | 50.0-2.56 | 20.0-1.91 | 30.0-2.00 |
| | (2.46-2.42) | (2.60-2.56) | (1.94-1.91) | (2.05-2.00) |
| $R_{sym}$ (%) | 9.1 (61.9) | 5.9 (13.1) | 5.1 (64.2) | 8.2 (51.4) |
| $I / \sigma I$ | 14.8 (2.7) | 35.3 (19.3) | 21.4 (2.03) | 24.0 (1.65) |
| Completeness (%) | 99.9 (99.1) | 99.7 (100.0) | 97.1 (82.2) | 98.1 (82.2) |
| Redundancy | 4.2 (3.3) | 10.4 (10.0) | 3.3 (2.9) | 5.1 (2.7) |
| **Refinement** | | | | |
| Resolution (Å) | 19.94-2.42 | | 19.75-1.91 | 29.98-2.00 |
| No. reflections | 65,618 | | 49,195 | 38,285 |
| $R_{work}$ / $R_{free}$ (%) | 21.82/25.47 | | 16.40/19.39 | 18.72/24.72 |
| No. atoms | | | | |
| Protein | 12,246 | | 4,135 | 4,101 |
| Ligand/ion | 66 | | 136 | 65 |
| Water | 340 | | 287 | 136 |
| *B*-factors | | | | |
| Protein | 59.32 | | 44.37 | 48.85 |
| Ligand/ion | 51.41 | | 62.68 | 67.06 |
| Water | 51.40 | | 48.98 | 44.22 |
| R.m.s deviations | | | | |
| Bond lengths (Å) | 0.007 | | 0.004 | 0.017 |
| Bond angles (°) | 1.026 | | 0.833 | 1.894 |

*Values in parentheses are for highest-resolution shell. For positioning of ligands, see **Supplementary Fig. 14.**

**Supplementary Table** 3*: In vitro* activity data

**Figure 4b, left panel:** *Tth* MAG lipase (wt) activity towards MAG, DAG, TAG substrates [U mg$^{-1}$]

| MAG (C8) | **DAG (C8)** | **TAG (C8)** | **MAG (C18)** |
|---|---|---|---|
| 16.40 ± 1.30 | 0.23 ± 0.11 | 0.00 ± 0.30 | 0.79 ± 0.53 |

**Figure 4b, right panel:** *Tth* MAG lipase variant activity towards MAG substrate [U mg$^{-1}$]

| | **C8** |
|---|---|
| wt | 17.99 ± 1.00 |
| E43A | 5.97 ± 0.14 |
| E43K | 0.00 ± 0.27 |
| E72R | 0.00 ± 0.25 |
| Y154A | 13.49 ± 0.72 |
| Y154R | 5.32 ± 0.44 |
| S113A | 0.62 ± 0.28 |
| ΔLid | 0.13 ± 0.29 |

**Figure 4c:** *Tth* MAG lipase (wt) activity towards *p*NP substrate [U mg$^{-1}$]

| C2 | **C4** | **C5** | **C6** | **C8** | **C10** |
|---|---|---|---|---|---|
| 1.23 ± 0.06 | 3.46 ± 0.15 | 3.56 ± 0.24 | 2.77 ± 0.18 | 2.49 ± 0.16 | 1.67 ± 0.13 |
| C12 | C14 | C16 | C18 | ferulate | |
| 0.98 ± 0.05 | 0.55 ± 0.04 | 0.04 ± 0.01 | 0.01 ± 0.01 | 0.03 ± 0.03 | |

**Supplementary Table 4: Quantification of CD data**

| | Helix | Sheet | Turn | Sum | Other | RMSD[a)] | NRMSD[a)] |
|---|---|---|---|---|---|---|---|
| **wt** | 36.5 | 18.1 | 17.6 | 72.2 | 27.7 | 7.9 | 2.9 |
| **wt (PMSF)** | 35.5 | 18.9 | 17.4 | 71.8 | 28.2 | 8.2 | 3.1 |
| **E43A** | 36.5 | 18.1 | 17.6 | 72.2 | 27.7 | 7.9 | 2.9 |
| **E43K** | 35.5 | 18.9 | 17.4 | 71.8 | 28.2 | 8.2 | 3.1 |
| **E72R** | 37.2 | 17.6 | 17.0 | 71.8 | 28.2 | 7.3 | 2.6 |
| **S113A** | 36.6 | 18.1 | 17.5 | 72.2 | 27.8 | 8.3 | 3.1 |
| **Δ(lid)** | 38.2 | 16.9 | 17.0 | 72.1 | 28.0 | 9.8 | 3.5 |
| **Y154A** | 38.4 | 17.1 | 16.8 | 72.3 | 27.7 | 7.0 | 2.5 |
| **Y154R** | 39.2 | 16.5 | 16.8 | 72.5 | 27.5 | 7.3 | 2.5 |

[a)] Root mean squares deviation and normalized root mean squares deviation as defined in CONTIN [3].

**Supplementary Table 5:** Water solubility of *p*-nitrophenyl FA esters used for *Tth* MAG lipase activity measurements

| Name | Corresponding acid | Water solubility (mg $ml^{-1}$)[1] |
|---|---|---|
| 4-Nitrophenyl acetate (*p*NP-C2) | Acetic/ethanoic acid | 1160 |
| 4-Nitrophenyl butyrate (*p*NP-C4) | Butyric/Butanoic acid | 24 |
| 4-Nitrophenyl valerate (*p*NP-C5) | Valeric/Pentanoic acid | 9 |
| 4-Nitrophenyl hexanoate (*p*NP-C6) | Caproic/Hexanoic acid | 15 |
| 4-Nitrophenyl octanoate (*p*NP-C8) | Caprylic/Octanoic acid | 1.513 |
| 4-Nitrophenyl decanoate (*p*NP-C10) | Capric/Decanoic acid | 0.1515 |
| 4-Nitrophenyl laurate (*p*NP-C12) | Lauric/Dodecanoic acid | 0.01504 |
| 4-Nitrophenyl myristate (*p*NP-C14) | Myristic/Tetradecanoic acid | 0.001481 |
| 4-Nitrophenyl palmitate (*p*NP-C16) | Palmitic/Hexadecanoic acid | 0.0001449 |
| 4-Nitrophenyl stearate (*p*NP-C18) | Stearic/Octadecanoic acid | 0.00001409 |

**Supplementary Table 6:** Steady-state kinetics of *Tth* MAG lipase variants with *p*-nitrophenyl substrates of different chain lengths

| ***p*NP-C2** | $K_M$ [mM] | $k_{cat}$ [$s^{-1}$] | $k_{cat}/K_M$ [$M^{-1}$ $s^{-1}$] | $(k_{cat}/K_M)_{variant/wt}$ |
|---|---|---|---|---|
| Wt | 2.3 ± 0.3 | 5.2 ± 0.6 | 2.2 ± 0.4 x $10^3$ | 1.0 |
| E43A | 1.9 ± 0.3 | 1.6 ± 0.2 x $10^1$ | 8.3 ± 1.4 x $10^3$ | 3.7 |
| E43K | 6.3 ± 0.6 x $10^{-1}$ | 1.0 ± 0.0 | 1.5 ± 0.2 x $10^3$ | 0.7 |
| E72R | 7.1 ± 0.7 x $10^{-1}$ | 4.0 ± 0.2 | 5.7 ± 0.6 x $10^3$ | 2.5 |
| Y154A | 1.4 ± 1.1 x $10^{-1}$ | 2.1 ± 1.5 x $10^1$ | 1.5 ± 1.6 x $10^3$ | 0.7 |
| Y154R | 2.0 ± 0.3 | 1.3 ± 0.1 x $10^1$ | 6.3 ± 1.2 x $10^3$ | 2.8 |

| ***p*NP-C4** | $K_M$ [mM] | $k_{cat}$ [$s^{-1}$] | $k_{cat}/K_M$ [$M^{-1}$ $s^{-1}$] | $(k_{cat}/K_M)_{variant/wt}$ |
|---|---|---|---|---|
| wt | 7.1 ± 0.7 x $10^{-1}$ | 8.6 ± 0.5 | 1.2 ± 0.1 x $10^4$ | 1.0 |
| E43A | 6.7 ± 1.9 x $10^{-1}$ | 1.6 ± 0.2 x $10^1$ | 2.4 ± 0.8 x $10^4$ | 2.0 |
| E43K | 8.2 ± 0.8 x $10^{-1}$ | 4.4 ± 0.2 | 5.3 ± 0.6 x $10^3$ | 0.4 |
| E72R | 1.4 ± 2.0 | 5.6 ± 5.2 | 3.9 ± 6.5 x $10^3$ | 0.3 |
| Y154A | 3.4 ± 0.6 | 1.3 ± 0.2 x $10^1$ | 3.8 ± 0.9 x $10^3$ | 0.3 |
| Y154R | 7.2 ± 1.9 x $10^{-1}$ | 1.2 ± 0.2 x $10^1$ | 1.6 ± 0.5 x $10^4$ | 1.3 |

| ***p*NP-C5** | $K_M$ [mM] | $k_{cat}$ [$s^{-1}$] | $k_{cat}/K_M$ [$M^{-1}$ $s^{-1}$] | $(k_{cat}/K_M)_{variant/wt}$ |
|---|---|---|---|---|
| wt | 8.3 ± 1.3 x $10^{-1}$ | 8.4 ± 0.7 | 1.0 ± 0.2 x $10^4$ | 1.0 |
| E43A | 6.5 ± 1.6 x $10^{-1}$ | 1.5 ± 0.2 x $10^1$ | 2.2 ± 0.6 x $10^4$ | 2.2 |
| E43K | 4.5 ± 0.8 x $10^{-1}$ | 3.2 ± 0.3 | 7.3 ± 1.5 x $10^3$ | 0.7 |
| E72R | 9.4 ± 3.3 x $10^{-1}$ | 1.1 ± 0.2 x $10^1$ | 1.2 ± 0.5 x $10^4$ | 1.2 |
| Y154A | 1.4 ± 0.2 | 1.1 ± 0.1 x $10^1$ | 8.1 ± 1.6 x $10^3$ | 0.8 |
| Y154R | 8.8 ± 2.2 x $10^{-1}$ | 1.3 ± 0.2 x $10^1$ | 1.5 ± 0.4 x $10^4$ | 1.5 |

| ***p*NP-C6** | $K_M$ [mM] | $k_{cat}$ [$s^{-1}$] | $k_{cat}/K_M$ [$M^{-1}$ $s^{-1}$] | $(k_{cat}/K_M)_{variant/wt}$ |
|---|---|---|---|---|
| wt | 3.6 ± 0.8 x $10^{-1}$ | 4.9 ± 0.4 | 1.4 ± 0.3 x $10^4$ | 1.0 |
| E43A | 7.1 ± 3.0 x $10^{-1}$ | 1.2 ± 0.3 x $10^1$ | 1.8 ± 0.8 x $10^4$ | 1.3 |
| E43K | 3.2 ± 0.9 x $10^{-1}$ | 2.8 ± 0.3 | 8.7 ± 2.7 x $10^3$ | 0.6 |
| E72R | 6.3 ± 3.4 x $10^{-1}$ | 8.3 ± 2.3 | 1.3 ± 0.8 x $10^4$ | 1.0 |
| Y154A | 4.8 ± 0.8 x $10^{-1}$ | 5.7 ± 0.4 | 1.2 ± 0.2 x $10^4$ | 0.9 |
| Y154R | 7.5 ± 3.6 x $10^{-1}$ | 1.2 ± 0.3 x $10^1$ | 1.5 ± 0.8 x $10^4$ | 1.1 |

| ***p*NP-C8** | $K_M$ [mM] | $k_{cat}$ [$s^{-1}$] | $k_{cat}/K_M$ [$M^{-1}$ $s^{-1}$] | $(k_{cat}/K_M)_{variant/wt}$ |
|---|---|---|---|---|
| wt | $4.9 \pm 1.5 \times 10^{-1}$ | $5.5 \pm 0.8$ | $1.1 \pm 0.4 \times 10^{4}$ | 1.0 |
| E43A | $5.2 \pm 3.4 \times 10^{-1}$ | $1.1 \pm 0.3 \times 10^{1}$ | $2.2 \pm 1.6 \times 10^{4}$ | 1.9 |
| E43K | $4.6 \pm 1.3 \times 10^{-1}$ | $3.5 \pm 0.5$ | $7.7 \pm 2.4 \times 10^{3}$ | 0.7 |
| E72R | $5.1 \pm 2.2 \times 10^{-1}$ | $6.4 \pm 1.3$ | $1.3 \pm 0.6 \times 10^{4}$ | 1.1 |
| Y154A | $6.2 \pm 2.3 \times 10^{-1}$ | $6.4 \pm 1.2$ | $1.0 \pm 0.4 \times 10^{4}$ | 0.9 |
| Y154R | $8.2 \pm 4.4 \times 10^{-1}$ | $1.1 \pm 0.3 \times 10^{1}$ | $1.3 \pm 0.8 \times 10^{4}$ | 1.2 |

| ***p*NP-C10** | $K_M$ [mM] | $k_{cat}$ [$s^{-1}$] | $k_{cat}/K_M$ [$M^{-1}$ $s^{-1}$] | $(k_{cat}/K_M)_{variant/wt}$ |
|---|---|---|---|---|
| wt | $3.0 \pm 0.7 \times 10^{-1}$ | $4.4 \pm 0.4$ | $1.5 \pm 0.4 \times 10^{4}$ | 1.0 |
| E43A | $2.7 \pm 1.0 \times 10^{-1}$ | $7.2 \pm 1.0$ | $2.\underline{6} \pm 1.0 \times 10^{4}$ | 1.8 |
| E43K | $3.3 \pm 1.1 \times 10^{-1}$ | $2.8 \pm 0.4$ | $8.6 \pm 2.9 \times 10^{3}$ | 0.6 |
| E72R | $1.3 \pm 0.7 \times 10^{-1}$ | $2.4 \pm 0.4$ | $1.9 \pm 1.1 \times 10^{4}$ | 1.3 |
| Y154A | $2.2 \pm 0.6 \times 10^{-1}$ | $2.9 \pm 0.3$ | $1.3 \pm 0.4 \times 10^{4}$ | 0.9 |
| Y154R | $3.9 \pm 1.1 \times 10^{-1}$ | $7.7 \pm 0.9$ | $2.0 \pm 0.6 \times 10^{4}$ | 1.3 |

| ***p*NP-C12** | $K_M$ [mM] | $k_{cat}$ [$s^{-1}$] | $k_{cat}/K_M$ [$M^{-1}$ $s^{-1}$] | $(k_{cat}/K_M)_{variant/wt}$ |
|---|---|---|---|---|
| wt | $1.1 \pm 0.1 \times 10^{-1}$ | $1.9 \pm 0.1$ | $1.8 \pm 0.2 \times 10^{4}$ | 1.0 |
| E43A | $1.0 \pm 0.1 \times 10^{-1}$ | $3.2 \pm 0.1$ | $3.3 \pm 0.5 \times 10^{4}$ | 1.8 |
| E43K | $5.3 \pm 1.3 \times 10^{-2}$ | $9.4 \pm 0.6 \times 10^{-1}$ | $1.8 \pm 0.5 \times 10^{4}$ | 1.0 |
| E72R | $1.7 \pm 0.6 \times 10^{-2}$ | $4.0 \pm 0.3 \times 10^{-1}$ | $2.4 \pm 0.9 \times 10^{4}$ | 1.3 |
| Y154A | $6.4 \pm 0.9 \times 10^{-2}$ | $1.1 \pm 0.0$ | $1.7 \pm 0.2 \times 10^{4}$ | 0.9 |
| Y154R | $1.2 \pm 0.2 \times 10^{-1}$ | $2.8 \pm 0.1$ | $2.4 \pm 0.3 \times 10^{4}$ | 1.3 |

| ***p*NP-C14** | $K_M$ [mM] | $k_{cat}$ [$s^{-1}$] | $k_{cat}/K_M$ [$M^{-1}$ $s^{-1}$] | $(k_{cat}/K_M)_{variant/wt}$ |
|---|---|---|---|---|
| wt | $1.2 \pm 0.1 \times 10^{-1}$ | $5.0 \pm 0.1 \times 10^{-1}$ | $4.2 \pm 0.4 \times 10^{3}$ | 1.0 |
| E43A | $1.0 \pm 0.1 \times 10^{-1}$ | $8.3 \pm 0.2 \times 10^{-1}$ | $8.2 \pm 0.8 \times 10^{3}$ | 1.9 |
| E43K | $4.3 \pm 1.2 \times 10^{-2}$ | $1.7 \pm 0.1 \times 10^{-1}$ | $3.9 \pm 1.1 \times 10^{3}$ | 0.9 |
| E72R | $4.4 \pm 1.4 \times 10^{-2}$ | $3.4 \pm 0.3 \times 10^{-1}$ | $7.7 \pm 2.6 \times 10^{3}$ | 1.8 |
| Y154A | $9.0 \pm 1.3 \times 10^{-2}$ | $3.8 \pm 0.2 \times 10^{-1}$ | $4.2 \pm 0.6 \times 10^{3}$ | 1.0 |
| Y154R | $1.7 \pm 0.7 \times 10^{-1}$ | $1.0 \pm 0.1$ | $6.1 \pm 2.6 \times 10^{3}$ | 1.5 |

#### Supplementary Table 7: FAME data

##### Normalized activity (ratio variant *Tth* lipase/ wt *Tth* lipase)

| | wt | ΔLid | S113A | E43A | E43K | Y154A | Y154R |
|---|---|---|---|---|---|---|---|
| | 1.080 | 0.616 | 0.680 | 2.148 | 0.962 | 0.983 | 1.595 |
| | 0.992 | 0.894 | 0.704 | 2.091 | 0.902 | 0.915 | 1.686 |
| | 1.084 | 0.840 | 0.437 | 1.769 | 0.614 | 1.059 | 0.756 |
| | 1.046 | 0.498 | | | 0.618 | 0.894 | |
| | 0.918 | 0.952 | | | | | |
| | 0.879 | 0.644 | | | | | |
| | | 0.447 | | | | | |
| No. experiments | n=6 | n=7 | n=3 | n=3 | n=4 | n=4 | n=3 |

##### Statistical analysis: unpaired t-test; two-tailed p values (GraphPad Prism 5.04)

| | wt | ΔLid | S113A | E43A | E43K | Y154A | Y154R |
|---|---|---|---|---|---|---|---|
| Mean | 1.000 | 0.699 | 0.607 | 2.003 | 0.774 | 0.963 | 1.346 |
| SD | 0.086 | 0.198 | 0.148 | 0.204 | 0.184 | 0.075 | 0.513 |
| p value *vs*. wt | | 0.0055 | 0.0013 | < 0.0001 | 0.0288 | 0.4985 | 0.1285 |
| p value summary | | ** | ** | *** | * | | |

**Supplementary Table 8: *Tth* MAG lipase sequence/structure relations to other α/β hydrolases**

| Sequence ID | Protein name | Organism | Phylum | Reference |
|---|---|---|---|---|
| I9KU82 | MAG lipase | *T. thermohydrosulfuricus* | Firmicutes | |
| D3YEX6 | Cinnamoyl esterase | *L. johnsonii* | Firmicutes | 2 |
| D2YW37 | Feruloyl esterase | *B. proteoclasticus* | Firmicutes | 1 |
| D0MHY8 | Osmotically inducible protein C | *R. marinus* | Bacteroidetes | 4 |
| B9JYM4 | Hydroxymuconic semialdehyde hydrolase | *A. vitis* | Proteobacteria | unpublished |
| Q6PE15 | Palmitoyl-protein thioesterase A | *M. musculus* | (Vertrebrate) | 5 |
| Q83AV9 | Alpha/beta hydrolase | *C. burnetii* | Proteobacteria | 6 |
| O30361 | Xylan esterase | *T. saccharolyticum* | Firmicutes | unpublished |
| Q65EQ1 | Carboxylesterase | *B. licheniformis* | Firmicutes | unpublished |
| Q1QEU6 | Triacylglycerol lipase | *P. cryohalolentis* | Proteobacteria | unpublished |
| A0A6I8WFT7 | MAG lipase | *P. ferrophilus* | Euryarchaeota | unpublished |
| B0V9K7 | Dienelactone hydrolase | *A. baumannii* | Proteobacteria | unpublished |
| A0A6L8NUA5 | Bromoperoxidase | *B. anthracis* | Firmicutes | unpublished |
| P82597 | MAG lipase | *Bacillus sp.* | Firmicutes | 7 |
| O07427 | MAG lipase | *M. tuberculosis* | Actinobacteria | 8 |
| A0A0B5WSQ6 | Lysophospho- lipase | *B. coagulans* | Firmicutes | 9 |
| J4WAT9 | AB hydrolase-1 | *B. bassiana* | Ascomycota | 10 |

| Sequence ID | Score[a] | E value[a] | PDB ID[b,c] | Q score[b] | Rmsd[b] | Seq. identity[b] | Lid[d] type | Lid length[d] | Assembly |
|---|---|---|---|---|---|---|---|---|---|
| I9KU82 | | | 7Q4J | | | | HBH | 43 | Dimer |
| D3YEX6 | 117.0 | 1.00E-31 | 3S2Z* | 0.72 | 1.18 | 0.32 | HBH | 46 | Dimer |
| D2YW37 | 102.0 | 1.00E-25 | 2WTM* | 0.68 | 1.31 | 0.30 | HBH | 46 | Dimer |
| D0MHY8 | 82.0 | 2.00E-17 | 5CML | 0.44 | 2.22 | 0.23 | HBH** | 48 | Dimer |
| B9JYM4 | 55.8 | 9.00E-09 | 3LLC | 0.56 | 1.88 | 0.22 | HBH | 50 | Monomer |
| Q6PE15 | 51.6 | 3.00E-07 | 6NY9 | 0.52 | 1.99 | 0.20 | HBH | 51 | Monomer |
| Q83AV9 | 47.4 | 4.00E-06 | *3TRD* | | | | *no* | | *Monomer* |

| | | | | | | | | | |
|---|---|---|---|---|---|---|---|---|---|
| O30361 | 39.3 | 0.003 | 3FCY | 0.32 | 2.11 | 0.20 | HB | 43 | Trimer |
| Q65EQ1 | 38.5 | 0.005 | 6NKG | | | | HB | 54 | Monomer |
| Q1QEU6 | 37.4 | 0.014 | 4NS4 | 0.36 | 2.03 | 0.22 | HB | 54 | Monomer |
| A0A6I8WFT7 | 34.7 | 0.1 | 6QE2 | | | | HBH | 66 | Dimer |
| B0V9K7 | 34.7 | 0.089 | 4OPM | | | | HB | 70 | Dimer |
| A0A6L8NUA5 | 33.5 | 0.23 | 3FOB | | | | HB | 80 | Trimer |
| P82597 | 32.7 | 0.41 | 4KE8* | 0.45 | 2.16 | 0.21 | HBH | 51 | Dimer |
| O07427 | 30.4 | 2.30 | 6EIC | 0.45 | 1.86 | 0.18 | HBH | 70 | Trimer |
| A0A0B5WSQ6 | 28.9 | 7.7 | 5O7G* | 0.39 | 2.13 | 0.15 | HB | 88 | Monomer |
| J4WAT9 | | | 7D79* | 0.49 | 2.00 | 0.19 | HBH | 84 | Dimer |

A BLAST search using the *Tth* MAG lipase sequence (A0A1I1X7Z7) against UniProtKB restricted to sequences with known 3D structures revealed 106 hits. From this pool of sequences, 15 non-redundant validated targets were selected **(Supplementary Fig. 12)**. These sequences were further analyzed by structural superposition using the PDBeFold server [11]. For ten out of fifteen selected hits, PDBeFold produced a reliable structure-based alignment for both catalytic domain segments (**Supplementary Fig. 5**). In addition, an α/β hydrolase from *Beauveria bassiana* (J4WAT9) was included, which was detected in the PDBeFold search but not in the BLAST search **(Supplementary Data 1)**.

[a] Taken from **Supplementary Data 1.**

[b] Taken from PDBeFold output using the structure of *Tth* MAG lipase as template against the PDB.

[c], *, indicates PDB code cited represents multiple structures deposited in PDB of the same sequence.

[d], for further details see **Fig. 7a**; **, indicates a non-canonical HBH lid domain (for further details, see text)

#### Supplementary Table 9: Primers used for mutagenesis

| |
|---|
| **E43A** |
| Forward: 5′-GGTTTTACAGGCAATAAAGTA**GCG**TCTCACTTTATTTTTGTGAAG-3′<br>Reverse: 5′-CTTCACAAAAATAAAGTGAGACGCTACTTTATTGCCTGTAAAACC-3′ |
| **E43K** |
| Forward: 5′-GGTTTTACAGGCAATAAAGTA**AAG**TCTCACTTTATTTTTGTGAAG-3′<br>Reverse: 5′-CTTCACAAAAATAAAGTGAGACTTTACTTTATTGCCTGTAAAACC-3′ |
| **E72R** |
| Forward: 5′-GACTTTTATGGTTCTGGA**AGA**AGTGATGGGGACTTTAGTGAA-3′<br>Reverse: 5′-TTCACTAAAGTCCCCATCACTTCTTCCAGAACCATAAAAGTC-3′ |
| **S113A** |
| Forward: 5'-AGAATAGGACTACTTGGTTTG**GCC**ATGGGAGGAGCTATTGCAGGG-3'<br>Reverse: 5'-CCCTGCAATAGCTCCTCCCATGGCCAAACCAAGTAGTCCTATTCT-3' |
| **Y154R** |
| Forward: 5′-AACGAAAGTGTAAAGCAA**CGC**GGAGCTATTATGGAA-3′<br>Reverse: 5′-TTCCATAATAGCTCCGCGTTGCTTTACACTTTCGTT-3′ |
| **Y154A** |
| Forward: 5′-AACGAAAGTGTAAAGCAA**GCC**GGAGCTATTATGGAA-3′<br>Reverse: 5′-TTCCATAATAGCTCCGGCTTGCTTTACACTTTCGTT-3′ |
| **Lid deletion mutant (Δ140-183)** |
| Forward: 5′-AAAAGGTACCTTTGAGCTGTCAAAAGGATACGAT-3′<br>Reverse: 5′-AAAAGGTACCAAAAGCTGGAGCCCATAGCACCA-3′ |
| **Subcloning from pETBlue-1 to pETM-14 for WT and mutants (SLiCe cloning)** |
| Forward: 5′-CTGGAAGTTCTGTTCCAGGGGCCCATGCAAAAGGCTGTTGAAATT – 3′<br>Reverse: 5′-CTTGTCGACGGAGCTCGAATTCGGCTATCCCTTTAACAATTCCTT-3′ |

**Supplementary Table 10: *Tth* MAG lipase structure lysine methylation sites**

| Residue | 7Q4J(A) | 7Q4J(B) | | 7Q4H(A) | 7Q4H(B) |
|---|---|---|---|---|---|
| **3** | | D | | | D |
| **12** | D | | | M | |
| **25** | | D | | | M |
| **27** | | | | | M |
| **41** | | | | | |
| **50** | | | | | |
| **57** | | D | | | |
| **94** | D | M | | | |
| **97** | D | | | | |
| **127** | D | D | | | |
| **131** | | M | | | D |
| **152** | | M | | | |
| **170** | D | D | | | |
| **173** | | | | | |
| **181** | | | | | |
| **189** | | | | M | |
| **193** | | D | | D | |
| **194** | M | M | | | D |
| **209** | M | D | | | |
| **216** | | | | | |
| **236** | D | D | | D | M |
| **242** | M | | | | |
| **243** | D | D | | | |
| **253** | M | M | | M | D |
| **254** | | M | | | |
| **258** | | | | | |
| **Methyl groups (total)** | **18** | **24** | | **7** | **11** |

M, mono-methylated; D, di-methylated.

**Supplementary Table 11: Geometry of C18 MAG-Ser113 intermediate (7Q4J)**

| ***Tth* MAG lipase chains** | **A** | **B** |
|---|---|---|
| Distances [Å] | | |
| C1(STE)-C2(STE) | 1.53 | 1.56 |
| C1(STE)-O1(STE) | 1.33 | 1.34 |
| C1(STE)-OG(Ser113) | 2.18 | 2.30 |
| C1(STE)-O1(GOL) | 2.60 | 2.59 |
| Angles [degrees] | | |
| C2(STE)-C1(STE)-O1(STE) | 110.9 | 112.6 |
| C2(STE)-C1(STE)-OG(Ser113) | 140.0 | 112.3 |
| C2(STE)-C1(STE)-O1(GOL) | 142.9 | 172.3 |
| O1(STE)-C1(STE)-OG(Ser113) | 95.3 | 89.1 |
| O1(STE)-C1(STE)-O1(GOL) | 71.7 | 74.6 |
| OG(Ser113)-C1(STE)-O1(GOL) | 73.1 | 69.4 |

Atom definitions are taken from PDB entry 7Q4J.

#### Supplementary Source Data

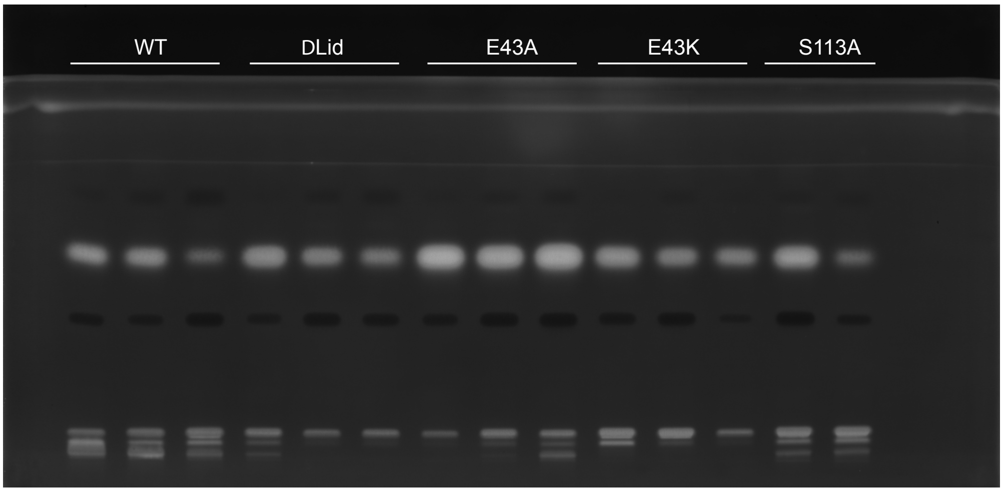

Uncropped scan of Figure 4f

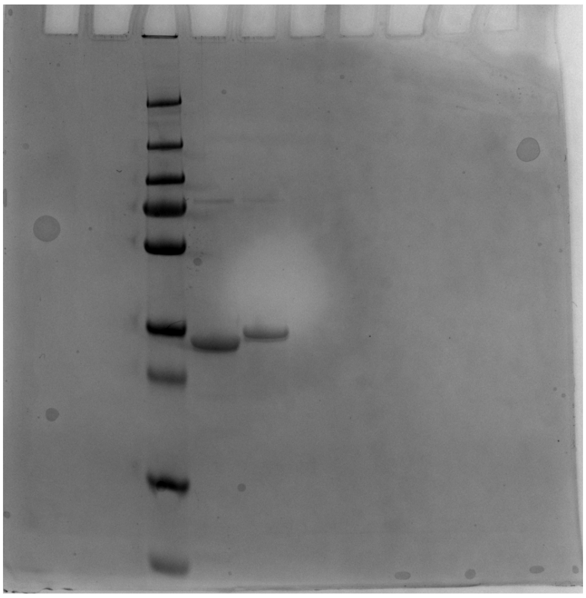

Uncropped scan of Supplementary Figure 13
